# Estimating occupancy dynamics and encounter rates with species misclassification: a semi-supervised individual-level approach

**DOI:** 10.1101/2021.03.17.433917

**Authors:** Anna I. Spiers, J. Andrew Royle, Christa L. Torrens, Maxwell B. Joseph

**Affiliations:** Earth Lab, Cooperative Institute for Research in Environmental Sciences, University of Colorado Boulder, CO, USA; Department of Ecology and Evolutionary Biology, University of Colorado Boulder, CO, USA; US Geological Survey Eastern Ecological Science Center; Institute of Arctic and Alpine Research, University of Colorado, Boulder, CO, USA

**Keywords:** carabid, imperfect classifier, morphospecies, NEON, observation error, occupancy models, semi-supervised, species misclassification

## Abstract

1. Large-scale, long-term biodiversity monitoring is essential to meeting conservation and land management goals and identifying threats to biodiversity. However, multispecies surveys are prone to various types of observation error, including false positive/negative detection, and misclassification, where a species is encountered but its species identity is not correctly identified. Previous methods assume an imperfect classifier produces species-level classifications, but in practice, particularly with human observers, we may end up with extraspecific classifications including “unknown”, morphospecies designations, and taxonomic identifications coarser than species. Disregarding these types of species misclassification in biodiversity monitoring datasets can bias estimates of ecologically important quantities such as demographic rates, occurrence, and species richness.
2. Here we develop an occupancy model that accounts for species non-detection and misclassification. Our framework accommodates extinction and colonization dynamics, allows for additional uncertain ‘morphospecies’ designations in the imperfect species classifications, and makes use of individual specimen with known species identities in a semi-supervised setting. We compare the performance of our joint classification-occupancy model to a reduced classification model that discards information about occupancy and encounter rate on a withheld test set. We illustrate our model with an empirical case study of the carabid beetle (Carabidae) community at the National Ecological Observatory Network Niwot Ridge Mountain Research Station, west of Boulder, CO, USA, and quantify taxonomist identification error by accounting for classification probabilities.
3. Species occupancy varied through time and across sites and species. The model yielded high probabilities (30 to 92% medians) of classification where the imperfect classifier matched the true species. The classification model informed by occupancy and encounter rates outperformed the classification that was not, and these differences were most pronounced for abundant species.
4. Our probabilistic framework can be applied to datasets with imperfect species detection and classification. This model can identify commonly misclassified species, helping biodiversity monitoring organizations systematically prioritize which samples need validation by an expert. Our Bayesian approach propagates classification uncertainty to offer an alternative to making conservation decisions based on point estimates

## 1 Introduction

Large-scale, long-term biodiversity monitoring is essential to meeting conservation and land management goals and identifying threats to biodiversity. Such comprehensive datasets increasingly include multispecies surveys that capture information-rich co-occurrence data, enabling community-level analyses (Iknayan et al., 2014; Ovaskainen et al., 2017). However, multispecies surveys are prone to various types of imperfect detection, including false absences where a species is present but not detected (Dorazio and Royle, 2005), and misidentification, where a species is encountered but its species identity is not correctly recorded (Miller et al., 2011).

Occupancy models account for observation error in biodiversity surveys that seek to understand species distributions, track population changes, and describe mechanisms underlying population and community dynamics (MacKenzie et al., 2002). Latent presence/absence states are modeled explicitly, with an observation model that accounts for the details of the detection process, including the potential for false negatives (non-detections at occupied sites) and false positives (detections at unoccupied sites) (Royle and Link, 2006; Miller et al., 2012; Chambert et al., 2015; Wright et al., 2020). Disregarding false positives in biodiversity monitoring datasets can bias estimates of ecologically important quantities such as demographic rates, occurrence, and species richness (McClintock et al., 2010; Chambert et al., 2015, 2018).

Multi-species surveys are also subject to errors in species identifications by imperfect classifiers. Imperfect classifiers include citizen scientists (e.g., North American Breeding Bird Survey (Sauer et al., 2017)), technicians trained in local taxonomy (e.g., invertebrate trapping by NEON (Hoekman et al., 2017)), automated methods (e.g., bat acoustic recording software (Wright et al., 2020) or convolutional neural networks used with camera trap data (Tabak et al., 2019)). Previous methods assume an imperfect classifier produces species-level classifications, but in practice, particularly with human observers, we may end up with extraspecific classifications including “unknown”, morphospecies designations, and taxonomic identifications coarser than species.

If species are prone to misclassification, then samples with known species identities might be used to correct estimates of occupancy parameters. However, using these data presents a methodological challenge. We refer to this situation as “semi-supervised”: true species identities are known for some but not all individuals. Previous multi-species occupancy models that accommodate misclassification have used multinomial models that sum over all individuals (Wright et al., 2020), or site-level validation data where the occupancy state of a species is known only at a site- or plot-level but not at an individual-level (Chambert et al., 2018). Using individual-level validation data requires a different approach.

Misclassified species identities can be dealt with using one of two contrasting approaches. A simple two step approach 1) uses a classifier to assign species IDs to each individual (creating one complete synthetic dataset from classifier output, for which species identities are treated as known), then 2) analyzes the constructed dataset using a downstream model (e.g., an occupancy model). This two step approach does not propagate uncertainty in species identity to the downstream model, and the assignment of species identities in the first stage does not use any information about occupancy or encounter rates. In contrast, a joint model directly uses classifier output as data, relating the observation process to underlying ecological states in one step. Such an approach can simultaneously account for uncertainty in species identities, and leverage information about occupancy and encounter rates to inform species identity estimates (Wright et al., 2020). However, there remains the practical question of how much value is added by a joint model vs. a two-stage approach. *A priori*, we expect that a joint model should produce better estimates of true species identities by using information on occupancy and encounter rates, but this has not yet been tested.

Here we develop an individual-level, semi-supervised, dynamic occupancy model that accounts for species nondetection and misclassification. Our Bayesian approach propagates classification uncertainty to offer an alternative to making conservation decisions based on point estimates. Our framework extends the classification occupancy model of Wright et al. (2020) to 1) accommodate extinction and colonization dynamics, 2) allow for additional uncertain “morphospecies” designations in the imperfect species classifications, and 3) make use of labeled samples with known species identities in a semi-supervised setting. Further, we compare the performance of a classification occupancy model to a reduced classification model that discards information about occupancy and encounter rate on a withheld test set. We demonstrate our model with an empirical case study of the carabid beetle (Carabidae) community at the National Ecological Observatory Network (NEON) Niwot Ridge Mountain Research Station (NIWO), west of Boulder, CO, USA, and quantify taxonomist identification error by accounting for classification probabilities.

## 2 Materials and Methods

### 2.1 Modeling occupancy dynamics with misclassification

Consider data collected at sites *i* = 1,…, *N*, according to a robust design (Hoekman et al., 2017) where each site is visited *J* times within primary periods *t* = 1,…, *T*, where the occupancy states are assumed to be constant within primary periods.

#### 2.1.1 State model

We are interested in occupancy states and encounter rates for species *k* = 1,…, *K*. Sites are either occupied (*z_i,k,t_* = 1) or not (*z_i,k,t_* = 0). We assume that the occupancy states arise as Bernoulli random variables:

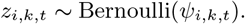

The probability of occupancy in the initial primary period is *ψ_i,k,1_*. Subsequent occupancy dynamics depend on the probability of colonization *γ_i,k,t_* and persistence *ϕ_i,k,t_*, such that for *t* > 1:

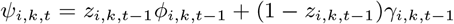

#### 2.1.2 Encounter model

On any particular sampling occasion *j*, we encounter *L_i,j,k,t_* individuals with encounter rate λ_*i,j,k,t*_. We assume that the number of encounters is a Poisson random variable: *L_i,j,k,t_* ~ Poisson(*z_i,k,t_*λ_*i,j,k,t*_). In a setting with misclassification, the number of encountered individuals *L_i,j,k,t_* is not observed directly because of uncertainty in the true species identities of encountered individuals. We do however observe the total number of individuals encountered on any particular occasion: 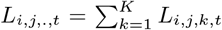. The properties of sums of Poisson random variables allow us to model these observed totals as:

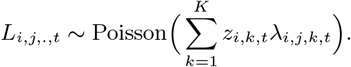

#### 2.1.3 Observation model

In addition to observing the total number of encountered individuals on an occasion *L_i,j,.,t_*, we assume that we also obtain imperfect species classifications for each encountered individual. In cases where individuals have been encountered (*L_i,j,.,t_* > 0), we obtain imperfect classifications of individuals *l* = 1,…, *L_i,j,., t_* and model these as arising from a categorical distribution with a species-specific probability vector:

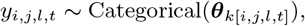

where *y_i,j,l,t_* is the imperfect classification, and ***θ***_*k*[*i,j,l,t*]_ is a probability vector associated with the true species of individual *l*, which we denote *k*[*i,j,l,t*]. Element *k*′ in the vector ***θ***_*k*[*i,j,l,t*]_ represents the probability that an individual is classified into category *k*′, conditional on the true species identity *k*[*i,j,l,t*], such that *θ*_*k*[*i,j,l,t*], *k*′_ = Pr(*y_i,j,l,t_* = *k*′ | *k*[*i,j,l,t*]). If species are always misclassified as other species, then *θ_k_* will be a vector of length *K* (Wright et al., 2020). If there are extraspecific classes (e.g. morphospecies), *θ_k_* may have more than *K* elements.

True species identities are modeled as:

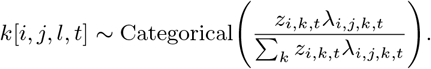

If ground truth species identity data are available for some individuals, then *k*[*i, j, l, t*] is partly observed and this model can be trained in a semi-supervised setting. In the unsupervised setting, this individual-level formulation is a disaggregated version of the single-season multinomial model of Wright et al. (2020) (Appendix S1).

#### 2.1.4 Incorporating morphospecies designations

In some settings the imperfect classifier might assign more classes than there are unique species, so that the vector ***θ**_k_* has more than *K* elements. For example, in the NEON beetle data, if a parataxonomist is unable to identify a set of similar individuals, they will classify those individuals as a unique morphospecies associated with that sampling occasion. Thus, it is possible for individuals to be classified into 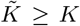 classes, where 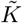 is sum of the number of species and the total number of morphospecies designations. In such cases, the matrix 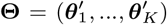 can be rectangular, with the first *K* columns corresponding to the classification probabilities for species 1,…, *K*, and the remaining columns corresponding to classification probabilities for non-species classes, e.g., morphospecies:

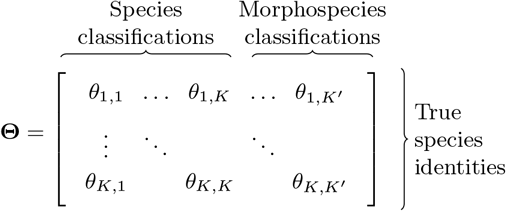

### 2.2 Case study

#### 2.2.1 Application to NEON carabid data

We fit our model to the carabid pitfall trap sampling data collected by NEON at NIWO in 2015-2018 (National Ecological Observatory Network, 2021). Carabids are a ubiquitous and speciose family of ground-dwelling invertebrates that are commonly collected by passive sampling methods, like pitfall traps, as described in Hoekman et al. (2017). Carabids are a well-studied sentinel group that make an excellent study system for assessing community occupancy rates and classification accuracy. Collecting and identifying carabids is resource-intensive, but NEON lowers this barrier to entry by providing a public carabid dataset with three levels of classification (parataxonomist, expert taxonomist, then DNA barcoding). Although NEON processes carabid samples at the domain level (sites within the same ecoregion) (Hoekman et al., 2017), we focus our analysis on one NEON site, NIWO, to assess occupancy across co-occurring species. We use the 2015-2018 dataset at NIWO since carabid sampling started in 2015 and expert classification data were not yet fully available for 2019 at the time of analysis in 2020 due to data latency (National Ecological Observatory Network, 2021). NIWO is a site in the southern Rocky Mountains, spanning subalpine conifer forest and alpine tundra.

We outline the relevant data collection protocol here, but Hoekman et al. (2017) offer more detail regarding NEON’s carabid pitfall trap data product. The sampling design at every NEON site consists of ten permanent plots across the site with four pitfall traps per plot. Traps are sampled and reset biweekly during the growing season, with a range of 5-7 collections per year at NIWO. Our model runs at the plot-level. In 2018 one plot was permanently relocated to ensure sampling was allocated proportionally to the NLCD cover types represented (NEON help desk, personal communication).

All carabid samples are classified by a parataxonomist, and a subset are sent to an expert taxonomist for validation (Figure 1) (Hoekman et al., 2017). Species classification by parataxonomists is considered imperfect due to the brief taxonomic training of parataxonomists. Identification by an expert taxonomist is treated as confirmation data and is limited due to budget constraints. We confirmed the accuracy of the expert taxonomist classifications in finding that all individuals sent for DNA barcoding by NEON match the expert taxonomist’s identification for the samples we used. In the few cases where the expert taxonomist could not identify a specimen to species-level, we use their genus-level classification for the validation dataset.

**Figure 1:**
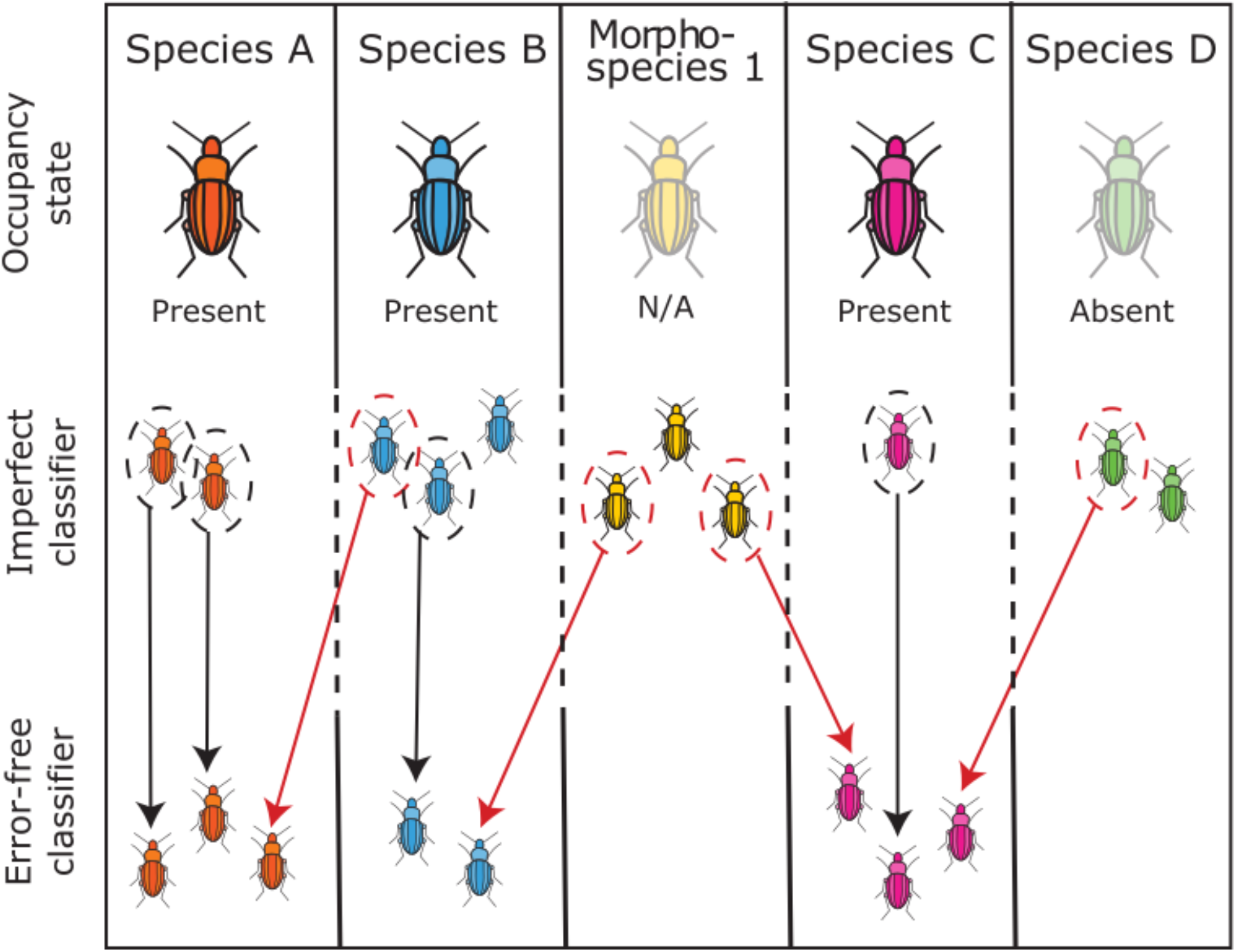
Classification scenarios in NEON carabid data. Each column is an imperfect classifier label. Each species is either present or absent, and morphospecies don’t have an occupancy state. In some cases, the imperfect classifier (parataxonomist) matches the error-free classifier (expert taxonomist) (black arrow), in other cases the imperfect classifier was wrong (red arrow), while in other cases still, the error-free classification is unknown due to lack of validation data. For example, the *Morphospecies 1* individual with no error-free classification must belong to a different column, but this species identity is unknown in the raw data.

Our dataset contains 4772 individuals, 1764 of which were identified by an expert taxonomist, and 62 species classified by the parataxonomist, 23 of which are morphospecies. Morphospecies identifications are unique to each NEON site and year. We fit our model using all individuals and used no environmental covariates. A hurdle for the NEON community in using the carabid pitfall trap data is reconciling parataxonomist and expert taxonomist classifications (Figure 1). Only one study to date has been published using the NEON carabid pitfall trap data (Egli et al., 2020), but Egli et al. (2020) analyze only the subset of individuals that have expert taxonomist classifications. Our model lowers the barrier of entry for using all imperfectly classified individuals while leveraging the available validation data.

#### 2.2.2 Model specification

We used informative priors for the species classification probability vectors ***θ***_1_,…, ***Θ***_*K*_ that placed higher probability density on the correct species classification. In the case of NEON beetle data, this is reasonable given the training that parataxonomists receive in beetle identification. Because all elements of each ***θ***_*k*_ vector need to sum to one, and each element is bounded between 0 and 1, we used a Dirichlet prior: ***θ***_*k*_ ~ Dirichlet(***α***_*k*_). We chose the Dirichlet concentration values ***α***_*k*_ by comparing draws from the Dirichlet prior distribution to our prior intuition about parataxonomist skill, using 200 along the diagonal (the element corresponding to the correct species identity), and 2 elsewhere.

We used multivariate normal priors at the species and site level, which allowed for correlations among parameters. These priors share information among initial occupancy, persistence, colonization, and encounter rates. The motivation for this stemmed from a prior expectation that these parameters could be related. For example, species that are more abundant might be more likely to occur initially, persist, or colonize new sites. Similar arguments could be made about relationships among parameters at a site level.

Each species is associated with a vector ***α***_*k*_ of length 4, where *α*_*k*,1_, *α*_*k*,2_, *α*_*k*,3_, and *α*_*k*,4_ are species-specific adjustments on initial occupancy, persistence, colonization, and encounter rates respectively. We assume that the species-specific adjustments are drawn from a multivariate normal prior with mean equal to zero, and an unknown covariance matrix: ***α***_*k*_ ~ Normal(**0, Σ**^(*α*)^). Similarly, site-specific adjustments **e**_i_ were drawn from a different multivariate normal prior. These adjustments were added together on a transformed scale to compute initial occupancy, persistence, colonization, and encounter rates, e.g., logit(*ψ*_*i,k*,1_) = *ϵ*_*i*,1_ + *α*_*k*,1_. A full model specification for the case study is available in Appendix S2 (Plummer et al., 2003).

To evaluate how the occupancy and encounter rate components of the full model informed classification probability estimates, we also developed a reduced model that discards all information about occupancy and abundance, using just the expert and para-taxonomist species classifications to estimate the classification matrix **Θ**. This comparison reveals the extent to which occupancy and encounter rates inform classification probabilities. If there are no differences in the estimates of classification probabilities, then a two-stage model which first models misclassification and then passes the posterior on as a prior for an occupancy/encounter model should perform as well as the joint model in which the classification model is integrated with the occupancy model. In addition to comparing posterior estimates for **Θ**, we withhold a randomly selected 20% of the imperfect classifications to evaluate which model (full or reduced) better predicts the data generated by the parataxonomist. All models were fit using JAGS (Appendix S3, dclone, and R v4.0.2 (Plummer et al., 2003; Sólymos, 2010; R Core Team, 2020) and visualized with ggplot (Wickham, 2016).

## 3 Results

### 3.1 Dynamic occupancy model

The occupancy model was designed to allow correlation between parameters across sites and species. Occupancy, growth, and turnover rates also varied through time. Sites with high encounter rates tended to have low initial occupancy and colonization probabilities and high persistence probabilities (Figure 2). Further, sites with high colonization rates tended to have high initial occupancy probabilities and low persistence probabilities. At the species level, we saw positive correlations among many of the model components, but in particular, species’ encounter rate was positively correlated with species’ initial occupancy, persistence, and colonization rates (Figure 2). Species varied in their detection success by the imperfect classifier, from ones that were common and consistently identified correctly (e.g. *Calathus advena*) to ones that were not identified at all (e.g., *Dicheirotrichus mannerheimii*) but were caught by the expert taxonomist.

**Figure 2:**
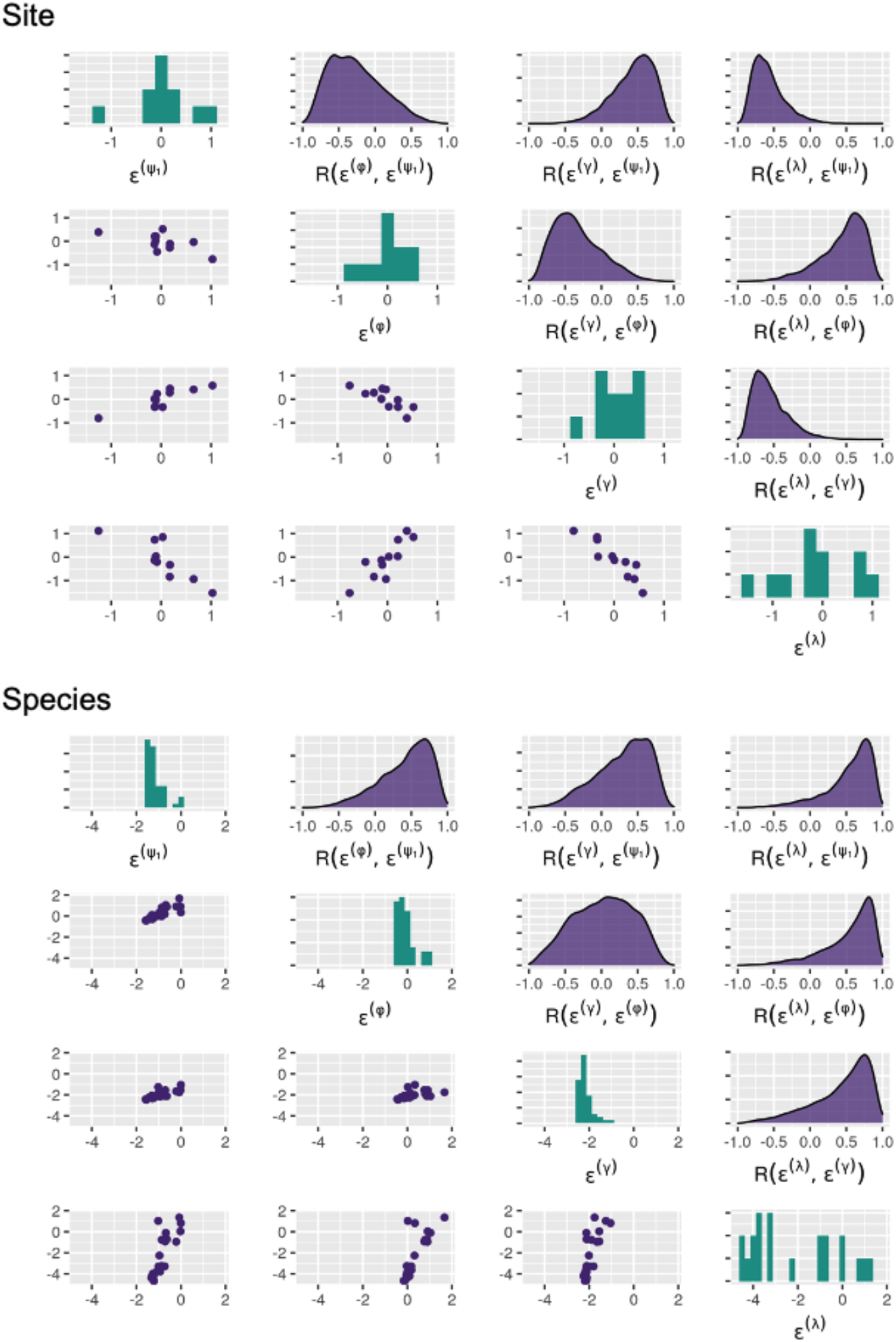
Random effects at the site and species levels. Rows correspond to the rows in the random effect covariance matrix: initial occupancy (row/col 1), persistence (row/col 2), colonization (row/col 3), and encounter rate (row/col 4). Along the diagonal are marginal histograms of posterior medians. Below the diagonal are pairwise scatterplots (each point is a site or species). Above the diagonal are posterior density plots of the pairwise correlation.

### 3.2 Classification model

The model yielded high probabilities of classification along the diagonal of the *θ* confusion matrix where the expert and para-taxonomist identifications match (Figure 3). The model favors the parataxonomist’s skill by giving more weight in the theta prior to diagonal values, making morphospecies classifications less probable. Individuals with morphospecies classifications make up a sizeable portion of the community, 811 out of the total 4772 total individuals identified by the parataxonomist. Despite the dirichlet priors favoring parataxonomist accuracy, some species had nontrivial probabilities of being classified as morphospecies than as the true species by the parataxonomist. For example, the parataxonomist was more likely to classify *Pterostichus* (*Hypherpes*) *sp*. as D13.2016.MorphBT than as the true species (Figure 3). However, no species had more than 3% probability (median) of being classified as another species (i.e., our model results indicate that the parataxonomist is most likely to identify a species either correctly or as a morphospecies).

**Figure 3:**
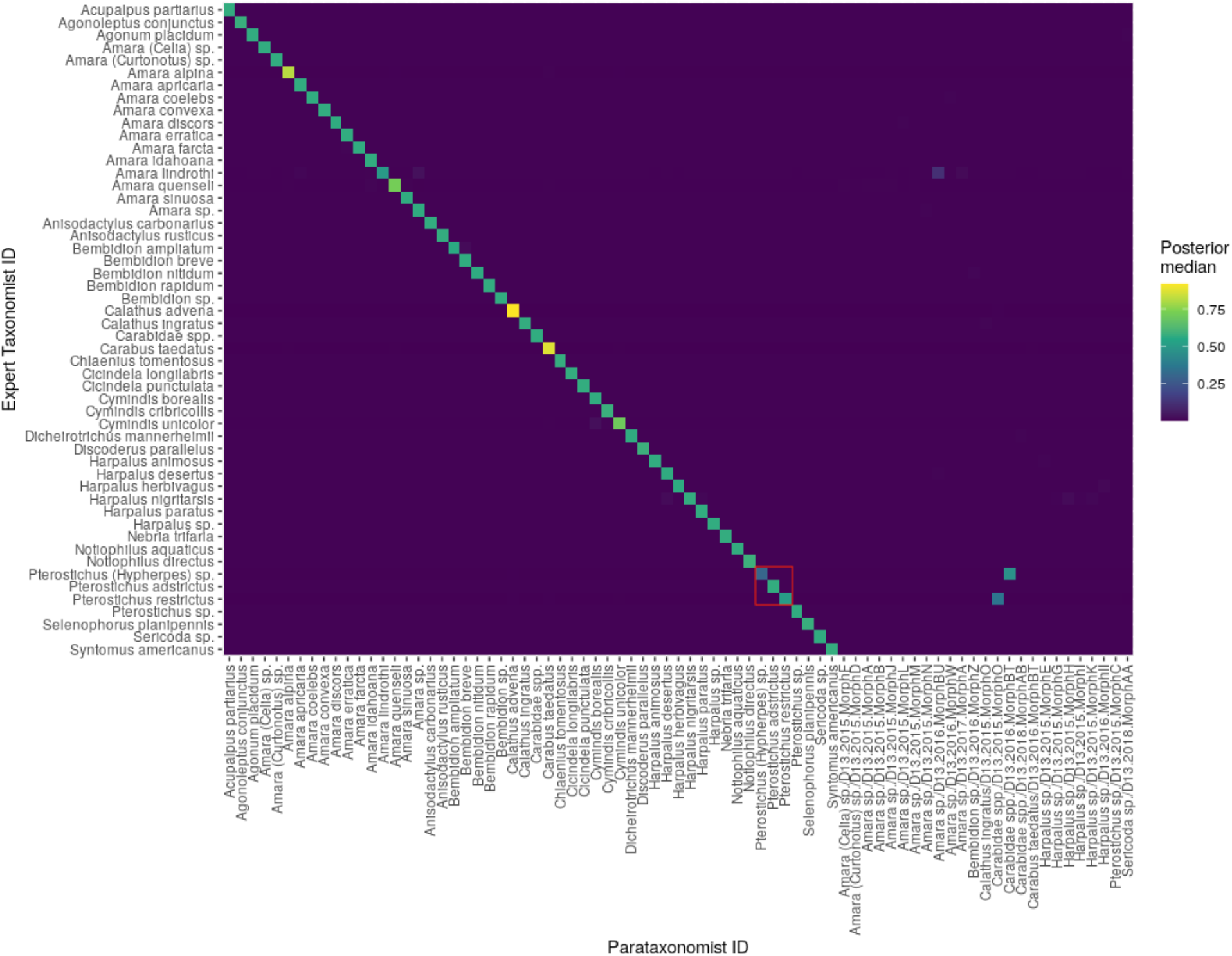
*θ* confusion matrix for the full model (classification with occupancy). Heat map of posterior median values where a value is interpreted as the probability that the species in that row is identified as a species in that column. Values along the diagonal are where the species is correctly identified. Values in each row sum to 1. The *θ* posterior distributions for the 3×3 cells outlined in red are illustrated in Figure 4, along with the posteriors for the reduced (classification alone) and prior models.

To evaluate the value added by informing the classification model with occupancy and encounter rates, we compared the full model to a reduced classification model that discards all information about occupancy and abundance. Most *θ_k_* probability vectors do not differ between the full and reduced model results. However, we see differences for a few species where there are non-overlapping *θ* posterior density distributions between the full and reduced models (i.e., Theta[46,46] and Theta [48,48], Figure 4). These differences are found most notably for the abundant species. The full model yielded higher classification probabilities for the abundant species. Further, the reduced model has wider 95% credible intervals compared to the full model for many theta indices (Figure 5). Thus, we find that a joint occupancy-classification model outperforms a two-stage model (classification, then occupancy).

**Figure 4:**
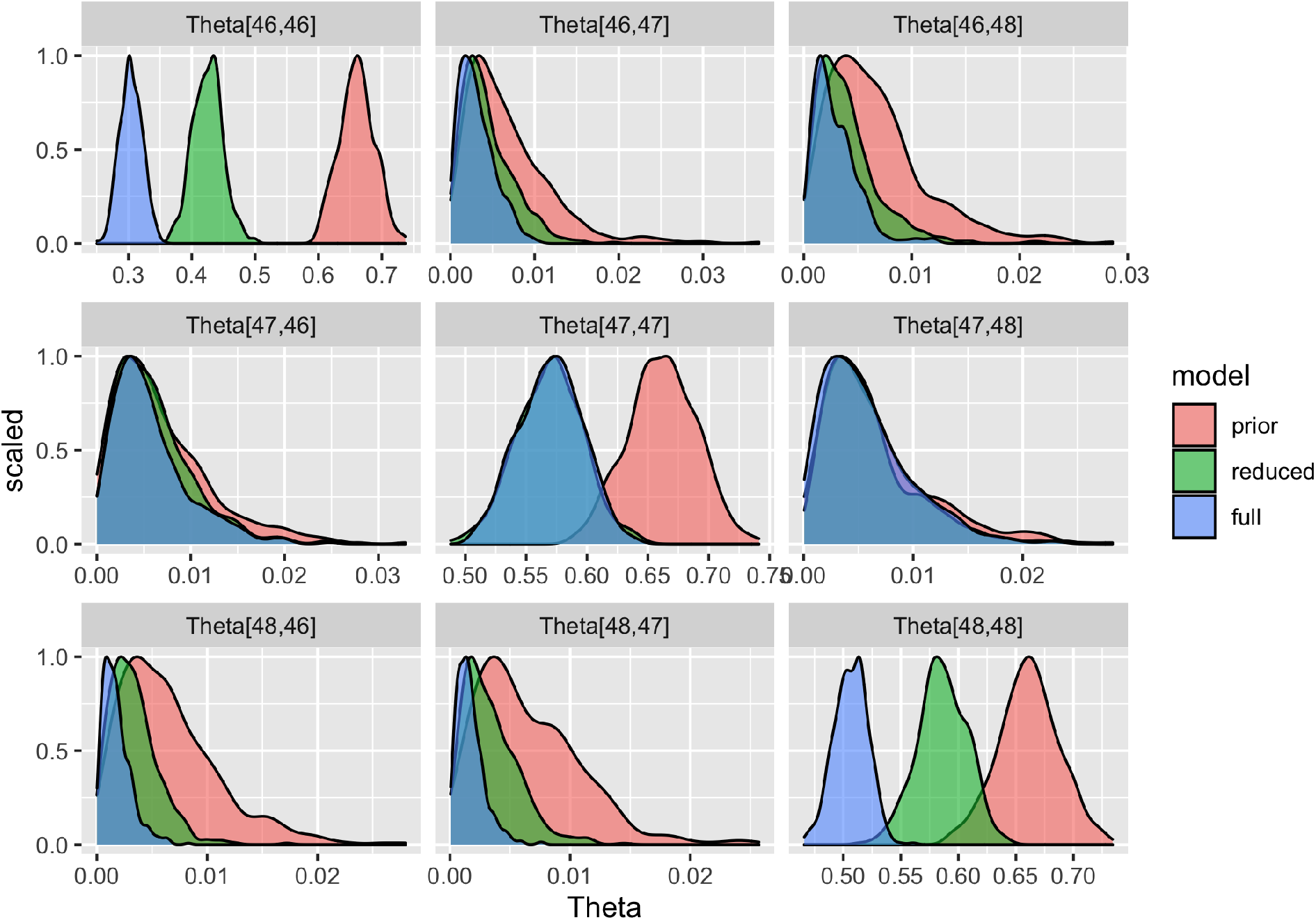
Comparison of *θ* density distribution between prior, full model posterior (classification with occupancy), and reduced model posterior (classification alone) for select *θ* confusion matrix indices. For non-abundant species, *θ_k_* probability distributions look like the second row. Here the three models’ probability distributions overlap on the off-diagonal (Theta[47,46]/[47,48]) and the posterior distributions overlap but differ from the prior along the diagonal (Theta[47,47]). In contrast the top and bottom rows reflect probability distributions for abundant species. The posterior distributions overlap and are narrower and smaller than the prior on off-diagonal values (Theta[46, 47]/[46,48], Theta[48,46]/[48,47]). Along the diagonal, we see a difference in posterior probability distribution between the reduced and full models (Theta[46,46], Theta[48,48]), visualizing how the full and reduced models perform differently.

We evaluated the performance of the full and reduced models by withholding a randomly selected 20% (352 individuals) of the imperfect classifications that have an expert identification (1764 individuals). For the withheld individuals, the validation metric macro-averages are listed in Table 1. For every validation metric, the full model yields better results than the reduced model. The validation metrics calculated to the species level highlighted substantial differences between the models for common species. This confirms the result that occupancy dynamics improve classification model performance compared to using classification data alone.

**Table 1:** Validation metric macro-averages

|  | Full model | Reduced model |
| --- | --- | --- |
| Accuracy | 0.65 | 0.56 |
| Precision | 0.23 | 0.19 |
| Recall | 0.37 | 0.35 |
| F1 score | 0.65 | 0.59 |
| Holdout log-likelihood | -84.7 | -110 |

## 4 Discussion

We developed a statistical approach to improve classification in multispecies datasets that leverages occupancy dynamics. Our probabilistic framework can be applied to datasets with imperfect species detection and also errors in classification of samples. This approach builds on recent work on classification in occupancy models (Devarajan et al., 2020, and references therein) by evaluating the advantage of a joint occupancy-classification model, allowing imperfect classifications to outnumber species, and leveraging individual-level confirmation data in a semi-supervised setting. While analyses targeting species richness may be shielded to a certain extent from imperfect classification (Egli et al., 2020), any population- or community-level analyses with taxonomic specificity require an understanding of classification uncertainty in the data. Whereas imperfect classifiers offer classification point-estimates, our model provides a vector of probabilities for every species.

This is the first model to consider how occupancy and encounter rates contribute to improving classification. We found that our joint occupancy-classification model outperformed a reduced model that disregarded occupancy dynamics in estimating imperfect classification (Figure 6, Table 1). When looking at the validation metrics at the species level, the joint model surpasses the reduced model even more for abundant species. In Figure 5, we see that the full model posteriors have smaller CI widths than the reduced model for many species, which visualizes the superior precision of the full model.

**Figure 5:**
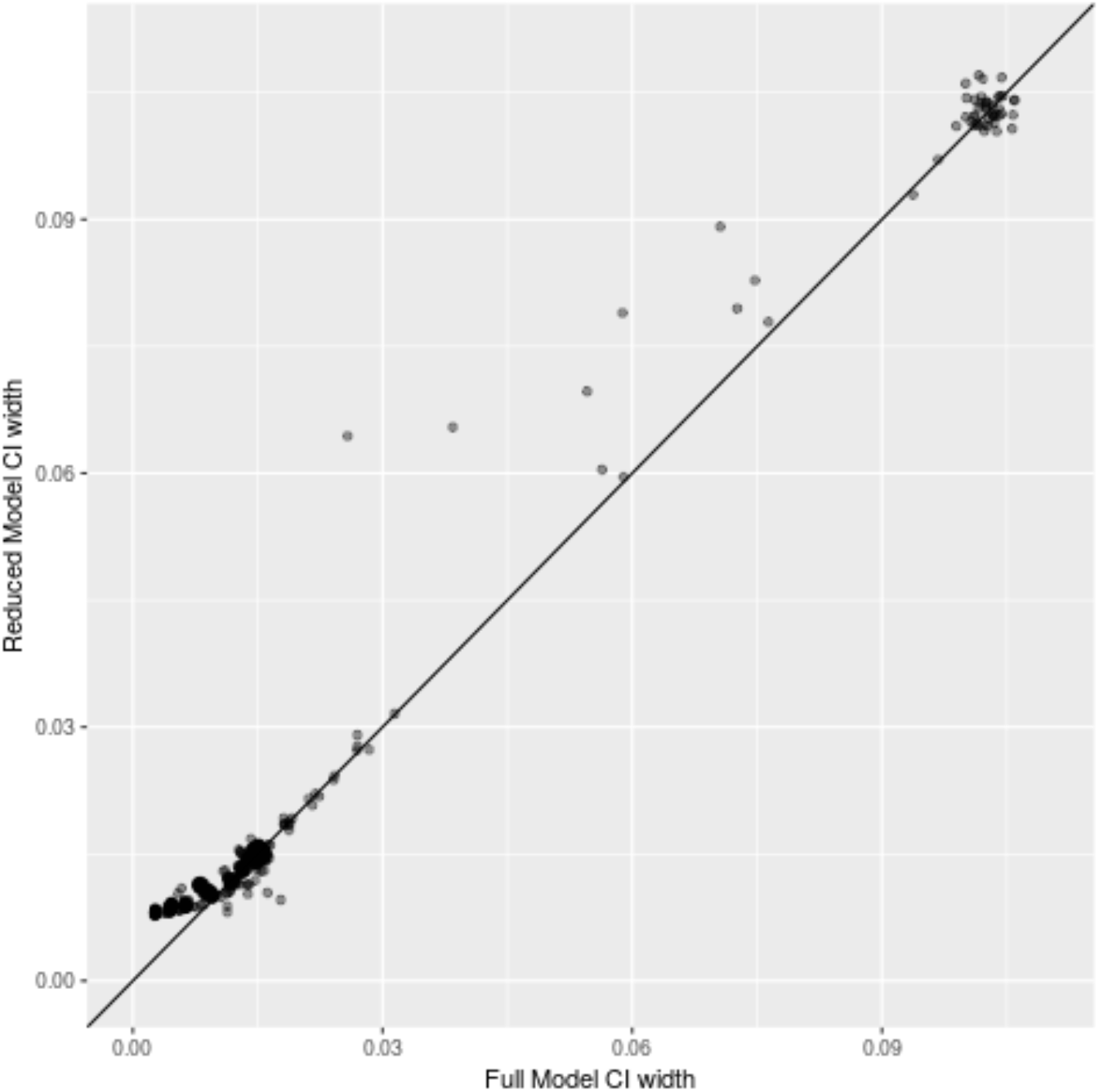
Comparison of precision between theta posterior 95% credible intervals (CI) of full model (occupancy with classification) and reduced model (classification alone). The full model is more precise for points above the line, indicating that the reduced model has a larger CI than the full model for that species.

**Figure 6:**
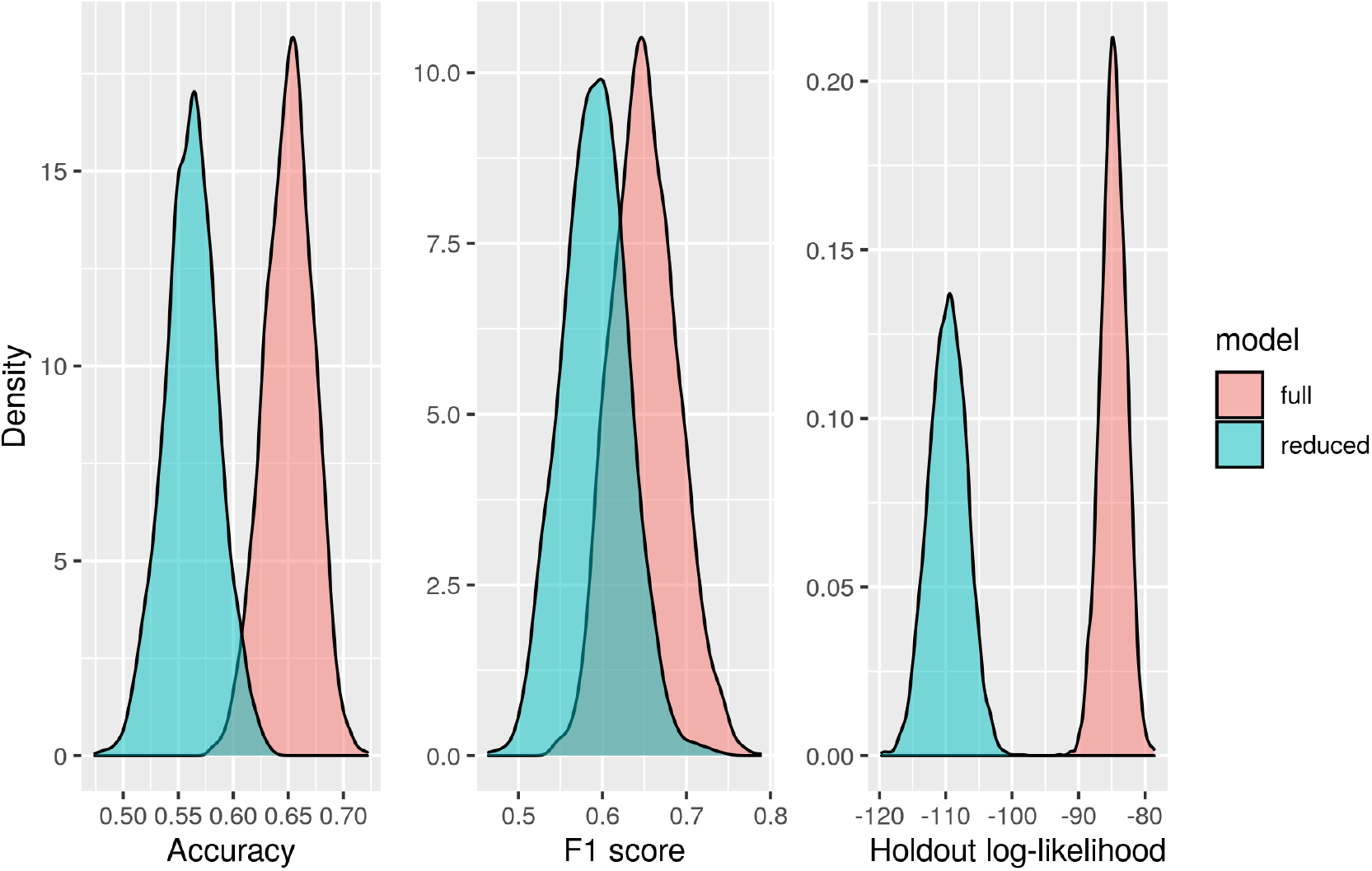
Validation metric macro-averages across posterior draws for accuracy, F1 score, and holdout log-likelihood.

False positive and negative species classifications are inevitable in any field collection, due to time and money constraints or imperfect classifiers (Royle and Link, 2006; Miller et al., 2012; McClintock et al., 2010; Hoekman et al., 2017). Accounting for false identifications is important to reduce bias in occupancy dynamics estimated from multispecies biodiversity monitoring datasets (McClintock et al., 2010; Miller et al., 2011; Chambert et al., 2015; Miller et al., 2015). Alternative models that account for false positives may consider data from only the focal species (Chambert et al., 2015) or from binary observations (Chambert et al., 2017). Like Wright et al. (2020), we use available counts from an imperfect classifier (Figure 1). However, we use all species detected, no matter how rare. By allowing taxonomic uncertainty propagation for multispecies datasets where imperfect classifications outnumber species (e.g., unknown, morphospecies, to the family level) by using a rectangular classification matrix *θ* (Figure 3), we remove an assumption that previous occupancy modeling methods have used (Chambert et al., 2018; Wright et al., 2020).

Our model is semi-supervised and makes use of data at the individual-level. Whereas alternative models use data pooled at the site- or occasion-level (Chambert et al., 2018; Wright et al., 2020), our model leverages the rich information at the individual-level to reveal which species are commonly mistaken by the imperfect classifier and how often. Individuals identified by the expert taxonomist we know as true positives so can be used as partially-observed occupancy data in our semi-supervised model. In contrast, the model by Wright et al. (2020) is unsupervised, but also at the individual-level. Because our analysis is done at the individual-level, we can use species counts to inform classification (Chambert et al., 2017). Although the model priors favor parataxonomist accuracy, the model found high probability of classification for a couple of morphospecies that were abundant in the data (D13.2015.MorphO and D13.2016.MorphBT) (Figure 3).

Despite its innovations, our model has limitations. For NEON’s carabid data from NIWO, which we used to fit our model, the parataxonomists were skilled and had high identification agreement with expert taxonomist classifications. The model may yield unexpected results when applied to a NEON site with lower parataxonomist accuracy. This raises the question of how much validation data is necessary to fit the model for varying degrees of imperfect classifier accuracy. This could be answered with simulations to identify what percentage or which type of samples should be prioritized for expert taxonomist classification to yield desired results.

We tried various iterations of the model before coming to the final disaggregated, semi-supervised, individual-level model. Using an aggregated data approach, we found the model either would not converge or struggled with identifiability, yielding multimodal posteriors for *θ*. Changing the Dirichlet priors to favor parataxonomist accuracy helped but did not eliminate the problem. Future work could more explicitly incorporate false positives by informing the *θ* priors with a list of species commonly misidentified by the imperfect classifier.

A model assumption is that samples were selected at random for verification. In reality, NEON prioritizes samples that were not classified to the species level by the parataxonomist. The model outlined here could be extended to represent processes such as this, by which samples are selected for verification by a more accurate classifier. Such extensions could take advantage of information contained in whether or not samples are selected for verification (e.g., the fact that a sample was not chosen for verification is informative, as it may indicate higher confidence in the initial classification).

Large-scale, long-term biodiversity surveys are critical to inform land management and conservation policy (Hughes et al., 2017) and require affordable and efficient species classifications to stay on track and within budget. Egli et al. (2020) make a case for better training of parataxonomists to improve classification error rates. However, training people is expensive and staff can be transient, calling for a more systematic solution. This probabilistic approach can model species occupancy while accounting for imperfect classification, without additional training. Innovations in occupancy models in general, are rapidly being made to consider an expanding variety of study systems and experimental designs (Bailey et al., 2014). Our results support the concept that ecological dynamics (i.e. occupancy and encounter rates) inform classification probabilities and lays a foundation for future work to build upon.

## Acknowledgements

The National Ecological Observatory Network is a program sponsored by the National Science Foundation and operated under cooperative agreement by Battelle Memorial Institute. This material is based in part upon work supported by the National Science Foundation through the NEON Program. We thank G. Vagle for taking part in the conception of this project and J. Coulombe for her graphical design assistance. The work was supported by the CU Boulder Grand Challenge investment in Earth Lab. AIS was supported as a GRA at Earth Lab for work on this project. Any use of trade, product, or firm names is for descriptive purposes only and does not imply endorsement by the U.S. Government.

## Authors’ contributions

AIS, CLT, and MBJ conceived the project idea; AIS, JAR, and MBJ designed methodology; AIS curated the data; AIS and MBJ analysed the data; AIS and MBJ led the writing of the manuscript. All authors contributed critically to the drafts and gave final approval for publication.

## Data availability

Carabid data accessible through the NEON data portal at https://data.neonscience.org/data-products/DP1.10022.001 (National Ecological Observatory Network, 2021). Code for data cleaning and analysis is available at https://github.com/annaspiers/NEON-NIWO-misclass

## Notes

https://github.com/annaspiers/NEON-NIWO-misclass

### Competing Interest Statement

The authors have declared no competing interest.

