## Supplemental appendices for "Estimating occupancy dynamics and encounter rates with species misclassification: a semi-supervised individual-level approach"

### Appendix S1: multinomial equivalence in the unsupervised setting

The disaggregated dynamic occupancy model that we present is equivalent to the multinomial model of (Wright et al., 2020) in a single-season, unsupervised setting. Fundamentally the equivalence is due to the link between a disaggregated categorical model, and an aggregated multinomial model.

We will outline the equivalence by first considering a set of observations made at one site $i$ and one occasion $j$, which consist of the total number of individuals encountered $L_{i,j,\cdot}$, and the imperfect species classifications for each encountered individual $y_{i,j,l}$ for $l = 1, ..., L_{i,j,\cdot}$. The disaggregated data model represents the probability of these observations as:

$$[L_{i,j,\cdot} \mid z_{i,1:K}, \lambda_{i,j,1:K}] \prod_{l=1}^{L_{i,j,\cdot}} [y_{i,j,l} \mid k_{i,j,l}, \theta][k_{i,j,l} \mid z_{i,1:K}, \lambda_{i,j,1:K}],$$

where $z_{i,1:K}$ is a vector of binary occupancy states for species $1, ..., K$, $k_{i,j,l}$ is the true species identity of individual $l$, and $\lambda_{i,j,1:K}$ is a vector of encounter rates.

In this appendix we will show that this is equivalent to the Poisson model of (Wright et al., 2020) (outlined in appendix S1 of their original paper), which consists of conditionally independent Poisson distributions for the number of individuals classified as each species $k' = 1, ..., K$:

$$[C_{i,j,\cdot,1}, ..., C_{i,j,\cdot,K} \mid z_{i,1:K}, \lambda_{i,j,1:K}, \theta] = \prod_{k'=1}^{K} \text{Poisson}\Big(C_{i,j,\cdot,k'} \mid \sum_{k=1}^{K} z_{i,k}\lambda_{i,j,k}\theta_{k,k'}\Big),$$

where $C_{i,j,\cdot,k'}$ is the total number of individuals at site $i$ on occasion $j$ classified as species $k'$, $z_{i,k}$ is the occupancy state of species $k$, $\lambda_{i,j,k}$ is the expected encounter rate, and $\theta_{k,k'}$ is the probability that an individual of species $k$ is classified as species $k'$. The proof of equivalence replaces the categorical distribution over class labels in the disaggregated model with a multinomial, then recovers conditionally independent Poisson distributions.

Consider one site and one occasion, dropping the subscripts $i$ and $j$ for simplicity. The model outlined above for the data from this site/occasion is:

$$[L_\cdot \mid z_{1:K}, \lambda_{1:K}] \prod_{l=1}^{L_\cdot} [y_l \mid k_l, \theta][k_l \mid z_{1:K}, \lambda_{1:K}],$$

where $L_\cdot$ is the total number of individuals encountered, $y_l$ is the (imperfect) classification of individual l, and $k_l$ is the true species identity of individual $l$.

The equivalence holds for the unsupervised case, where none of the true species identities are known. In this case we can marginalize over the unknown species identities as follows:

$$[L_\cdot \mid z_{1:K}, \lambda_{1:K}] \prod_{l=1}^{L_\cdot} \sum_{k_l=1}^{K} [y_l \mid k_l, \theta][k_l \mid z_{1:K}, \lambda_{1:K}],$$

Writing out the probability mass functions for the Poisson and categorical distributions:

$$= \frac{1}{L.!}\Big(\sum_{k=1}^{K} z_k\lambda_k\Big)^{L.} e^{-\sum_{k=1}^{K} z_k\lambda_k} \prod_{l=1}^{L.}\sum_{k_l=1}^{K} \frac{z_{k_l}\lambda_{k_l}\theta_{k_l,y_l}}{\sum_k z_k\lambda_k}$$

Note that the last term in this expression is the probability mass function for the observed classifications: $[y_l \mid \theta, z_{1:K}, \lambda_{1:K}] = \text{Categorical}\Big(\sum_{k_l=1}^{K} \frac{z_{k_l}\lambda_{k_l}\theta_{k_l,y_l}}{\sum_k z_k\lambda_k}\Big)$. Rewrite this categorical component as a multinomial, where $C_{.,k'}$ represents the total number of individuals classified as species $k'$.

$$= \frac{1}{L.!}\Big(\sum_{k=1}^{K} z_k\lambda_k\Big)^{L.} e^{-\sum_{k=1}^{K} z_k\lambda_k} \frac{L.!}{\prod_{k'} C_{.,k'}!}\Big(\sum_{k=1}^{K}\frac{z_k\lambda_k\theta_{k,1}}{\sum_k z_k\lambda_k}\Big)^{C_{.,1}} \times ... \times \Big(\sum_{k=1}^{K}\frac{z_k\lambda_k\theta_{k,K}}{\sum_k z_k\lambda_k}\Big)^{C_{.,K}}$$

Note that $L.!$ cancels out:

$$= \Big(\sum_{k=1}^{K} z_k\lambda_k\Big)^{L.} e^{-\sum_{k=1}^{K} z_k\lambda_k} \frac{1}{\prod_{k'} C_{.,k'}!}\Big(\sum_{k=1}^{K}\frac{z_k\lambda_k\theta_{k,1}}{\sum_k z_k\lambda_k}\Big)^{C_{.,1}} \times ... \times \Big(\sum_{k=1}^{K}\frac{z_k\lambda_k\theta_{k,K}}{\sum_k z_k\lambda_k}\Big)^{C_{.,K}}$$

Factor out the denominators in the mutinomial probabilities:

$$= \Big(\sum_{k=1}^{K} z_k\lambda_k\Big)^{L.} e^{-\sum_{k=1}^{K} z_k\lambda_k} \frac{1}{\prod_{k'} C_{.,k'}!}\Big(\frac{1}{\sum_k z_k\lambda_k}\Big)^{\sum_{k'} C_{.,k'}}\Big(\sum_{k=1}^{K} z_k\lambda_k\theta_{k,1}\Big)^{C_{.,1}} ... \Big(\sum_{k=1}^{K} z_k\lambda_k\theta_{k,1}\Big)^{C_{.,K}}$$

Now, recall that $L. = \sum_{k'} C_{.,k'}$, which leads to a convenient cancellation:

$$= \frac{\big(\sum_{k=1}^{K} z_k\lambda_k\big)^{L.}}{\big(\sum_{k=1}^{K} z_k\lambda_k\big)^{L.}} e^{-\sum_{k=1}^{K} z_k\lambda_k} \frac{1}{\prod_{k'} C_{.,k'}!}\Big(\sum_{k=1}^{K} z_k\lambda_k\theta_{k,1}\Big)^{C_{.,1}} ... \Big(\sum_{k=1}^{K} z_k\lambda_k\theta_{k,1}\Big)^{C_{.,K}}$$

$$= e^{-\sum_{k=1}^{K} z_k\lambda_k} \frac{1}{\prod_{k'} C_{.,k'}!}\Big(\sum_{k=1}^{K} z_k\lambda_k\theta_{k,1}\Big)^{C_{.,1}} ... \Big(\sum_{k=1}^{K} z_k\lambda_k\theta_{k,1}\Big)^{C_{.,K}}$$

Group like terms:

$$= e^{-z_1\lambda_1} ... e^{-z_K\lambda_K} \frac{1}{C_{.,1}!}\Big(\sum_{k=1}^{K} z_k\lambda_k\theta_{k,1}\Big)^{C_{.,1}} ... \frac{1}{C_{.,K}!}\Big(\sum_{k=1}^{K} z_k\lambda_k\theta_{k,1}\Big)^{C_{.,K}}$$

Then, note that $\exp(-z_k\lambda_k) = \exp(-\sum_{k'=1}^{K} z_k\lambda_k\theta_{k,k'})$:

$$= \prod_{k=1}^{K} e^{-\sum_{k'=1}^{K} z_k\lambda_k\theta_{k,k'}} \frac{1}{C_{.,1}!}\Big(\sum_{k=1}^{K} z_k\lambda_k\theta_{k,1}\Big)^{C_{.,1}} ... \frac{1}{C_{.,K}!}\Big(\sum_{k=1}^{K} z_k\lambda_k\theta_{k,1}\Big)^{C_{.,K}}$$

$$= \prod_{k=1}^{K}\prod_{k'=1}^{K} e^{-z_k\lambda_k\theta_{k,k'}} \frac{1}{C_{.,1}!}\Big(\sum_{k=1}^{K} z_k\lambda_k\theta_{k,1}\Big)^{C_{.,1}} ... \frac{1}{C_{.,K}!}\Big(\sum_{k=1}^{K} z_k\lambda_k\theta_{k,1}\Big)^{C_{.,K}}$$

Finally, we can group these terms to recover the conditionally independent Poisson random variables from (Wright et al., 2020):

$$= \Bigg(\frac{1}{C_{.,1}!}\Big(\sum_{k=1}^{K} z_k\lambda_k\theta_{k,1}\Big)^{C_{.,1}} e^{-\sum_{k=1}^{K} z_k\lambda_k\theta_{k,1}}\Bigg) ... \Bigg(\frac{1}{C_{.,K}!}\Big(\sum_{k=1}^{K} z_k\lambda_k\theta_{k,K}\Big)^{C_{.,K}} e^{-\sum_{k=1}^{K} z_k\lambda_k\theta_{k,K}}\Bigg)$$

$$= \prod_{k'=1}^{K} \text{Poisson}\Big(C_{\cdot,k'} \mid \sum_{k=1}^{K} z_k \lambda_k \theta_{k,k'}\Big),$$

which proves the equivalence between the disaggregated categorical observation model and the observation model described in (Wright et al., 2020).

### Appendix S2: case study model details

The NEON carabid beetle case study estimates occupancy dynamics and encounter rates for sites $i = 1, ..., N$, species $k = 1, ..., K$, and years $t = 1, ..., T$. The state and observation models are identical to those described in the main text. Occupancy dynamics were modeled as a first order Markov process. For the first year:

$$z_{i,k1} \sim \text{Bernoulli}(\psi_{i,k,1}).$$

In subsequent years:

$$z_{i,k,t} \sim \text{Bernoulli}(z_{i,k,t-1}\phi_{i,k,t-1} + (1 - z_{i,k,t-1})\gamma_{i,k,t-1}),$$

where $z_{i,k,t}$ is a binary occupancy state, $\phi_{i,k,t}$ is the probability of a site remaining occupied from year $t$ to year $t+1$, and $\gamma_{i,k,t}$ is the probability of an unoccupied site in year $t$ being occupied in year $t+1$.

We assume that encounter rates vary by site and species, leading to the following encounter model for occasion $j$:

$$L_{i,j,\cdot,t} \sim \text{Poisson}\Big(\sum_{k=1}^{K} z_{i,k,t}\lambda_{i,k}\Big).$$

The model for the observed species or morphospecies classifications is:

$$y_{i,j,l,t} \sim \text{Categorical}(\boldsymbol{\theta}_{k[i,j,l,t]}),$$

Classification probabilities were modeled using an informative Dirichlet prior:

$$\boldsymbol{\theta}_k \sim \text{Dirichlet}(\boldsymbol{\alpha}_k),$$

where $\alpha_{k,k}$ was set to 80, and all other elements $\alpha_{k,k'}$ such that $k \neq k'$ were set to 2. These specific values were chosen based on prior predictive simulations to match our expectations about the classification skill of parataxonomists.

The model for true species identities is:

$$k[i,j,l,t] \sim \text{Categorical}\left(\frac{z_{i,k,t}\lambda_{i,k}}{\sum_k z_{i,k,t}\lambda_{i,k}}\right).$$

We modeled initial occupancy ($\psi_{i,k,1}$), persistence ($\phi_{i,k}$), colonization ($\gamma_{i,k}$), and encounter rates ($\lambda_{i,k}$) as a function of site and species-specific random effects:

$$\text{logit}(\psi_{i,k,1}) = \epsilon_{i,1} + \alpha_{k,1},$$

$$\text{logit}(\phi_{i,k}) = \epsilon_{i,2} + \alpha_{k,2},$$

$$\text{logit}(\gamma_{i,k}) = \epsilon_{i,3} + \alpha_{k,3},$$

$$\log(\lambda_{i,k}) = \epsilon_{i,4} + \alpha_{k,4}.$$

These random effects were modeled with multivariate normal priors to allow for correlations among parameters. At the site level:

$$\begin{bmatrix} \epsilon_{i,1} \\ \epsilon_{i,2} \\ \epsilon_{i,3} \\ \epsilon_{i,4} \end{bmatrix} \sim \text{Normal}(\boldsymbol{\mu}, \boldsymbol{\Sigma}^{(\epsilon)}).$$

At the species level:

$$\begin{bmatrix} \alpha_{k,1} \\ \alpha_{k,2} \\ \alpha_{k,3} \\ \alpha_{k,4} \end{bmatrix} \sim \text{Normal}(\mathbf{0}, \boldsymbol{\Sigma}^{(\alpha)}).$$

The covariance matrices associated with these random effects were given inverse Wishart priors, with the parameters of the inverse Wishart distributions chosen based on prior predictive simulations:

$$\boldsymbol{\Sigma}^{(\epsilon)} \sim \text{Wishart}^{-1}(\boldsymbol{I}, 10),$$

$$\boldsymbol{\Sigma}^{(\alpha)} \sim \text{Wishart}^{-1}(\boldsymbol{I}, 10),$$

where $\boldsymbol{I}$ is a $4 \times 4$ identity matrix, and 10 represents the degrees of freedom parameter of the inverse Wishart distribution. Finally the vector of means, which acts essentially as an intercept parameter, was given a standard normal prior: $\boldsymbol{\mu} \sim \text{Normal}(\mathbf{0}, \boldsymbol{I})$. A graphical representation of the model is given in Figure S2.1.

We used simulation-based calibration (SBC) to check the validity of the sampler implemented by JAGS (Talts et al., 2018). Simulation-based calibration involves simulating many datasets from the prior predictive distribution of the model, and for each simulated dataset, drawing samples from the posterior. These posterior samples can then be compared to the true (known) values simulated from the prior in terms of rank statistics, which for well-behaved samplers are uniform. Non-uniform rank statistics are indicative of problems with the sampler, e.g., a tendency to under- or over-sample from the tails, or systematic directional bias (Talts et al., 2018). We simulated 3000 data sets from the prior distribution of the model in the two-species case, with 20 sites, 5 time steps, and 2 sampling occasions per site in each time step. Uniform SBC rank histograms indicate that the MCMC sampler built by JAGS is able to draw samples from the posterior distribution of the model (Fig. S2.2).

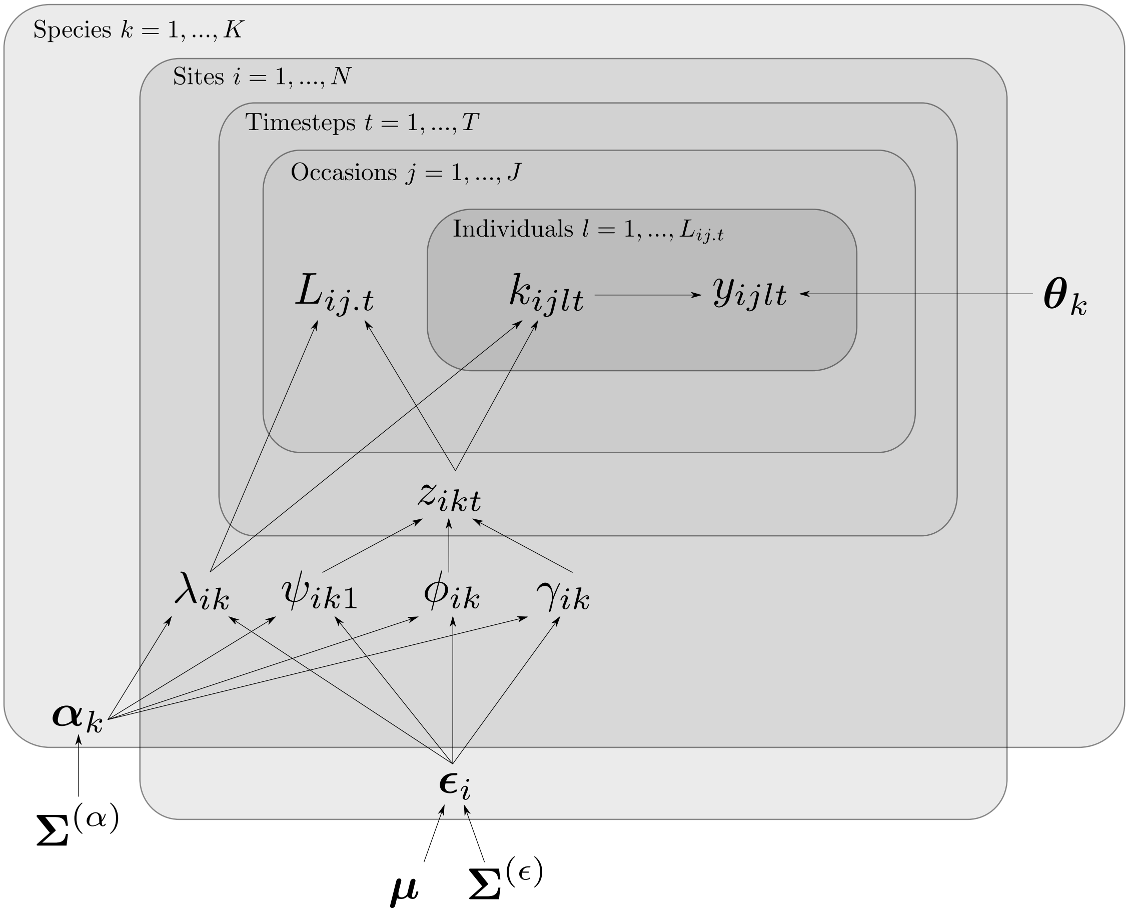


Figure S2.1: Directed acyclic graph representation of the NEON carabid beetle model. Grey plates represent levels of the model and edges represent dependence.

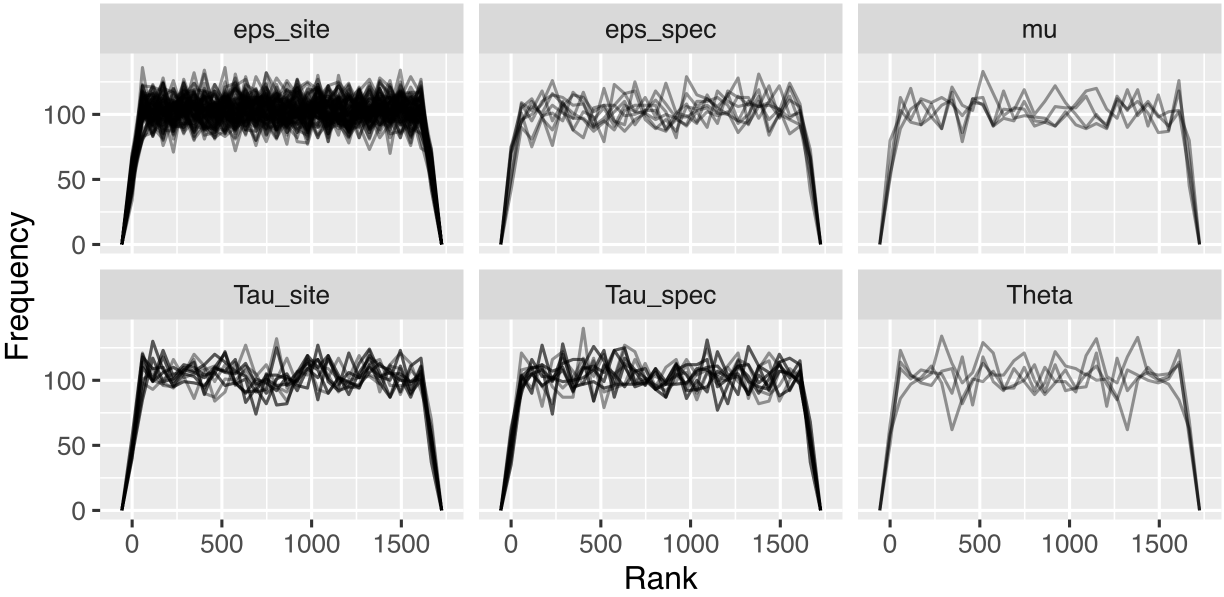


Figure S2.2: Simulation based calibration rank frequency polygons indicate uniformity. Each panel represents a different model parameter group, and each line represents a different parameter in that group (e.g., there are two species, so that the matrix $\mathbf{\Theta}$ is 2 by 2 so that there are four elements, and there are four lines in the Theta panel, one for each matrix element).

### Appendix S3: JAGS model specification

Also accessible in code repository (see Data availability statement).

```
model {
  # Priors
  for (i in 1:4) {
    mu_spec[i] ~ dnorm(0, 1)
    mu_site[i] = 0
  }
  Tau_spec[1:4, 1:4] ~ dwish(R[1:4, 1:4], 10)
  for ( i in 1:K_exp) {
    eps_spec[i, 1:4] ~ dmnorm(mu_spec, Tau_spec)
  }
  Tau_site[1:4, 1:4] ~ dwish(R[1:4, 1:4], 10)
  for (i in 1:nsite) {
    eps_site[i, 1:4] ~ dmnorm(mu_site, Tau_site)
  }
  for (i in 1:nsite){
    for (k in 1:K_exp) {
      logit_psi1[i, k] = eps_site[i, 1] + eps_spec[k, 1]
      psi1[i, k] = ilogit(logit_psi1[i, k])
      logit_phi[i, k] = eps_site[i, 2] + eps_spec[k, 2]
      phi[i, k] = ilogit(logit_phi[i, k])
      logit_gamma[i, k] = eps_site[i, 3] + eps_spec[k, 3]
      gamma[i, k] = ilogit(logit_gamma[i, k])
      log_lambda[i, k] = eps_site[i, 4] + eps_spec[k, 4]
      lambda[i, k] = exp(log_lambda[i, k])
    }
  }
  for (k in 1:K_exp) {
    Theta[k,1:K_para] ~ ddirch(alpha[k,1:K_para])
  }
  # State model
  for (i in 1:nsite) {
    for (k in 1:K_exp) {
      z[i, k, 1] ~ dbern(psi1[i, k])
      for (t in 2:nyear) {
        z[i, k, t] ~ dbern(z[i,k,t-1]*phi[i, k] +
                    (1 - z[i,k,t-1])*gamma[i, k])
      }
      for (t in 1:nyear) {
        zlam[i, k, t] = z[i, k, t] * lambda[i, k]
      }
    }
  }
  # Observation model
  for (i in 1:nsite) {
    for (j in 1:nsurv) {
      for (t in 1:nyear) {
          L[i, j, t] ~ dpois(sum(zlam[i, 1:K_exp, t]))
      }
    }
  }
  for (l in 1:Ltot) {
    pi[l, 1:K_exp] = zlam[site[l], 1:K_exp, year[l]] / sum(zlam[site[l], 1:K_exp, year[l]])
    k[l] ~ dcat(pi[l, 1:K_exp])
    y[l] ~ dcat(Theta[k[l], 1:K_para])
  }
  # Derived parameters
  for (i in 1:nsite) {
    for (k in 1:K_exp) {
      psi[i, k, 1] <- psi1[i, k]
      for (t in 2:nyear) {
        psi[i, k,t] = psi[i, k, t-1]*phi[i, k] +
                     (1 - psi[i, k, t-1])*gamma[i, k]
        log_growth[i, k, t] = log(psi[i, k, t]) - log(psi[i, k, t-1])
        turnover[i, k, t-1] = (1 - psi[i, k, t-1])*gamma[i, k]/psi[i, k, t]
      }
    }
  }
}
```
